# Chromatin loop dynamics during cellular differentiation are associated with changes to both anchor and internal regulatory features

**DOI:** 10.1101/2022.10.31.514600

**Authors:** Marielle L. Bond, Eric S. Davis, Ivana Y. Quiroga, Michael I. Love, Hyejung Won, Douglas H. Phanstiel

**Affiliations:** Curriculum in Genetics and Molecular Biology, University of North Carolina at Chapel Hill, NC, USA; Curriculum in Bioinformatics and Computational Biology, University of North Carolina at Chapel Hill, NC, USA; Thurston Arthritis Research Center, University of North Carolina, Chapel Hill, NC, USA; Department of Biostatistics, University of North Carolina, Chapel Hill, NC, USA; Department of Genetics, University of North Carolina, Chapel Hill, NC, USA; Lineberger Comprehensive Cancer Center, The University of North Carolina at Chapel Hill, Chapel Hill, North Carolina, USA; Neuroscience Center, The University of North Carolina at Chapel Hill, Chapel Hill, North Carolina, USA; Department of Cell Biology and Physiology, University of North Carolina, Chapel Hill, NC, USA

**Author notes:** Corresponding authors: Douglas H. Phanstiel and Hyejung Won.

## Abstract

3D chromatin structure has been shown to play a role in regulating gene transcription during biological transitions. While our understanding of loop formation and maintenance is rapidly improving, much less is known about the mechanisms driving changes in looping and the impact of differential looping on gene transcription. One limitation has been a lack of well powered differential looping data sets. To address this, we conducted a deeply sequenced Hi-C time course of megakaryocyte development comprising 4 biological replicates and 6 billion reads per time point. Statistical analysis revealed 1,503 differential loops. Gained loops were enriched for AP-1 occupancy and correlated with increased expression of genes at their anchors. Lost loops were characterized by increases in expression of genes within the loop boundaries. Linear modeling revealed that changes in histone H3 K27 acetylation, chromatin accessibility, and JUN binding in between the loop anchors were as predictive of changes in loop strength as changes to CTCF and/or cohesin occupancy at loop anchors. Finally, we built linear models and found that incorporating the dynamics of enhancer acetylation and loop strength increased accuracy of gene expression predictions.

## INTRODUCTION

The three dimensional (3D) organization of chromatin is thought to play an important role in transcriptional regulation and has been implicated in many biological processes, including cellular differentiation and response to external stimuli^1^. While several types of 3D chromatin structures exist, chromatin loops are of particular interest as they are thought to regulate gene expression by bringing distal regulatory elements (e.g. enhancers) into close physical proximity with gene promoters via point to point interactions. Indeed, loop anchors are typically enriched for enhancers and promoters and correlate with differences in gene expression^2–4^. Aberrations to chromatin looping are associated with a variety of human diseases and developmental disorders such as Cornelia de Lange syndrome^5^, polydactyly^6,7^, and cancer^8,9^. While the basic mechanisms of loop formation have been established, major questions remain regarding the mechanisms driving differential looping during biological development and their functional impact.

The vast majority of chromatin loops are thought to form through a process called loop extrusion, in which the cohesin complex is loaded onto DNA and reels in chromatin until it reaches convergently bound CTCF proteins^10^. In specific cell types and biological conditions chromatin loops can form through non-canonical mechanisms including phase separation^8,11,12^ or binding of lineage-specific factors like LDB1^8,11,12^. While chromatin loops have been shown to change over cellular transitions, the mechanisms that govern these structural changes, and the impact of these changes on gene expression, remain poorly understood^13,14^.

The relationship between looping and gene expression is even less clear. Cell type specific loops correlate with differential expression patterns, supporting a role for loops in gene regulation^15–17^. Moreover, forced looping between enhancers and promoters at select loci has been shown to activate transcription^18,19^. However, several recent studies have called the role of loops in regulating gene expression into question. Live cell imaging of looping between SOX2 and an enhancer known to regulate its expression showed no correlation between enhancer-promoter proximity and gene transcription^20^. In another study, rapid and thorough degradation of cohesin led to a complete removal of cohesin-driven loops in human cancer cells with only a minor impact on gene expression^14^. In summary, the degree to which chromatin looping regulates gene expression is still unresolved.

One impediment to answering these questions is that identifying differential loops between cells and conditions remains challenging. Due to the depth of sequencing required for Hi-C data sets, the statistical requirements of differential analysis, and the cost of DNA sequencing, most existing differential looping studies lack the statistical power to adequately identify differential loops. And without comprehensive and rigorously defined sets of differential loops, it is challenging to determine what mechanisms drive differential looping and what transcriptional impact they have.

To address this gap, we generated a deeply sequenced Hi-C data set characterizing the differentiation of K562 cells into a megakaryocyte-like state. By sequencing over 18 billion reads across three timepoints and four biological replicates, we achieved a statistical power of roughly 0.932 and identified 1,503 differential loops. Generation and intersection with accompanying maps of chromatin accessibility, histone acetylation, transcription factor (TF) binding, and gene expression revealed insights into both the mechanisms and the functional impacts of differential looping during cellular differentiation. Interestingly, we find that regulatory features both at and between loop anchors correlate with changes in loop strength. Finally, we show that incorporating H3 K27 acetylation and chromatin looping dynamics into linear models in addition to promoter acetylation improves predictions of changes in gene expression.

## RESULTS

### Differentiation of K562 cells induces large-scale changes to 3D chromatin structure across multiple scales

To understand how 3D chromatin structures change over cellular differentiation, we performed a deeply sequenced, 3 timepoint Hi-C time course tracking the differentiation of K562 cells into a megakaryocyte-like state^21^. We treated K562 cells with phorbol 12-myrisate 13-acetate (PMA), which has been shown to induce a megakaryocyte-like phenotype^21,22^, for 0, 6, and 72h. We confirmed differentiation using qPCR for *ITGB3*, a megakaryocyte marker^23^ (**Fig S1A**). As K562 cells differentiate into a megakaryocyte-like state, they lose their potential to differentiate into erythroid cells. We confirmed this with qPCR for *KLF1*, an erythroid marker, which decreases in expression over differentiation^24^ (**Fig S1B**). We then performed in situ Hi-C on four biological replicates and sequenced them to a depth of roughly 6 billion reads per time point (**Table S1**). We generated Hi-C contact maps using the Juicer pipeline^15^, identified compartments using the EigenVector package^25^, topologically associating domains (TADs) using arrowhead^15^, and chromatin loops using SIP^26^ (**Fig S1C**). Replicates exhibited high similarity as measured by principal component analysis (PCA) (**Fig S1D**).

Visual inspection of the data revealed clear changes at multiple scales including nuclear compartments, TADs, and chromatin loops (**Fig 1A-C**). To assess the sequencing depth and replicates required to achieve sufficient statistical power, we analyzed our data set using the RNASeqPower package^27^. Using our dispersion of 0.0019 and median sequencing depth of 38 counts per million (CPM) per loop, the statistical power to detect 2-fold changes was 0.932, which is generally considered to be well-powered^28^ (**Fig 1D**). We used the dispersion from our Hi-C data to model predicted statistical power across multiple different sequencing depths and numbers of replicates (**Fig 1D**). Holding sequencing depth per replicate constant, we found that decreasing to 3 or 2 replicates reduced the power estimates to 0.762 and 0.416 respectively. To determine how this increase in power translated into the number of differential loops, we subsampled our data to multiple depths and analyzed it with 2, 3, or all 4 replicates (**Fig 1E**). Using our full data set of four biological replicates (~900 million reads per replicate) we identified 1,503 differential loops using DESeq2^29^ with a log-2 fold change (LFC) greater than 1.5 (adjusted p-value < 0.05) compared to only 698 and 45 identified with 3 or 2 replicates respectively (**Fig 1E**). This underscores how critical sequencing depth is in the sensitivity to detect differential loops. Finally, we confirmed the quality of these differential loops through aggregate peak analysis^15^ (APA), showing that gained loops have higher contact at 72 vs 0 h, and lost loops have higher contact at 0 vs 72 h (**Fig 1F**).

**Figure 1.**
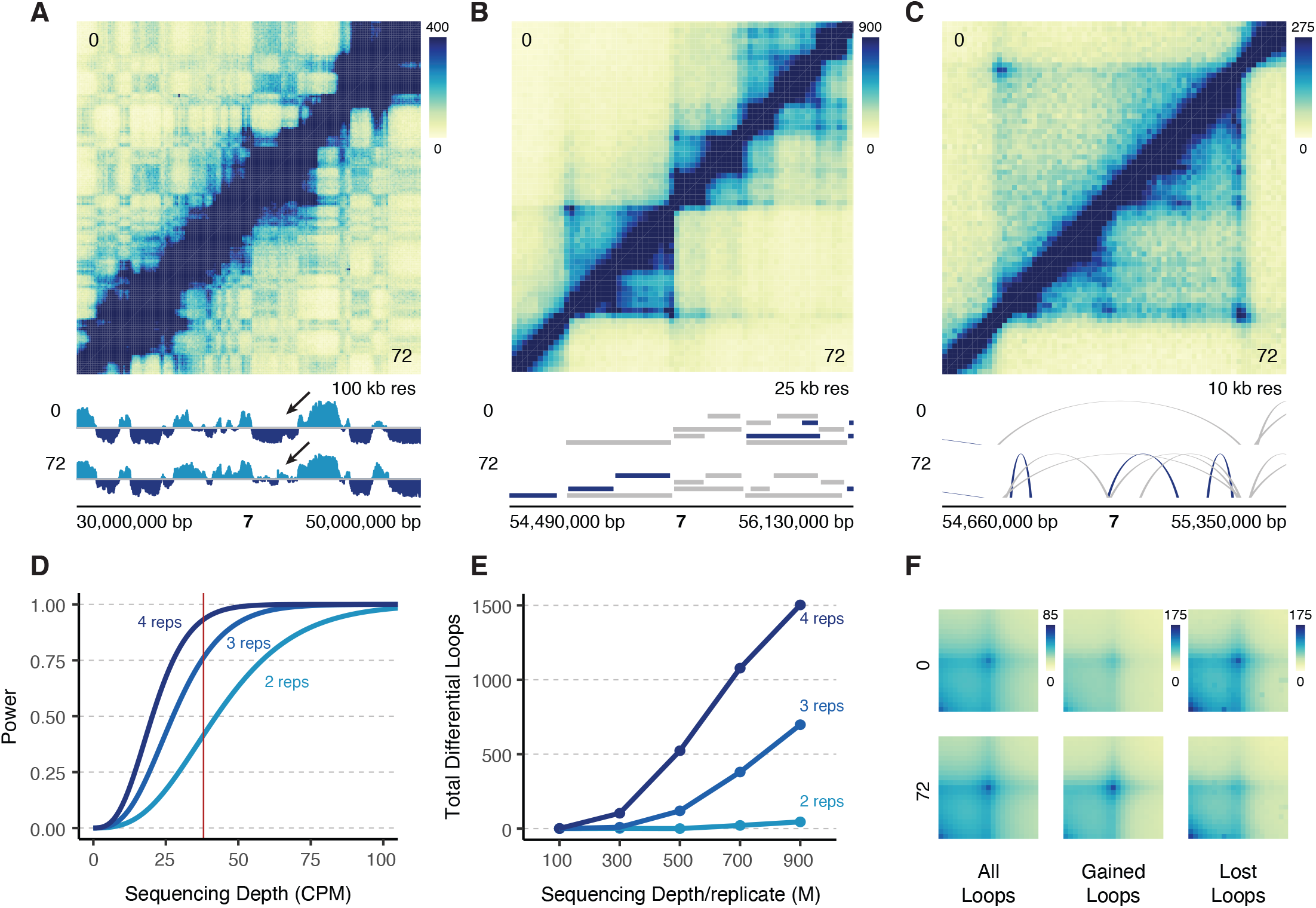
Deeply sequenced Hi-C experiments provide sensitive detection of differential chromatin loops. **(A)** 20 Mb region on chromosome 7 at 100kb resolution comparing K562s at 0h (top) to differentiated megakaryocytes at 72h (bottom). Signal tracks show compartmental eigenvector calls (light blue = compartment A, dark blue = compartment B). The arrow points to qualitative changes in compartmentalization. **(B)** Zoom in of a 1.6 Mb region of chromosome 7 at 25 kb resolution. TAD calls are indicated by ranges below the Hi-C map for each cell type (dark blue = cell type specific, gray = shared across cell types). **(C)** Zoom in of a 690 kb region in figure B at 10 kb resolution showing a region with differential loops. Arches indicate loop calls (dark blue = cell type specific, gray = shared across cell types). **(D)** Statistical power modeled across various theoretical sequencing depths at 2, 3, and 4 biological replicates. Red line indicates the median sequencing depth per loop (CPM: counts per million). **(E)** Actual number of differential loops called at multiple different subsampled sequencing depths for different numbers of replicates (M: millions). **(F)** Aggregate peak analysis for all loops, gained loops, and lost loops.

### Genes at the anchors of gained, but not lost, loops exhibit concordant changes in expression

To assess the potential transcriptional impacts of differential loops, we performed RNA-seq across 8 time points (0, 0.5, 1.5, 3, 6, 24, 48, and 72 h) in K562 cells treated with PMA. DESeq2 analysis identified 3,190 differential genes (adjusted p-value < 0.05, LFC > 2) which were grouped into 6 clusters using k-means clustering (**Fig 2A**). The biggest and most unique changes were observed at 6 and 72 hours (1,619 differential, 527 unique genes after 6 h; 1,925 differential, 236 unique genes after 72 h), which coincide exactly with our selected Hi-C timepoints. Consistent with the differentiation of these cells into a megakaryocyte-like state, the upregulated genes were enriched for Gene Ontology (GO) terms relating to cell differentiation, cell adhesion, and morphogenesis, and pathways including focal adhesion, hematopoietic cell lineage, and regulation of actin cytoskeleton (**Fig S2A-B**). Upregulated genes include megakaryocyte markers *VWF, FLI1*, and *ITGB3*, further supporting acquisition of a megakaryocyte-like phenotype.

**Figure 2.**
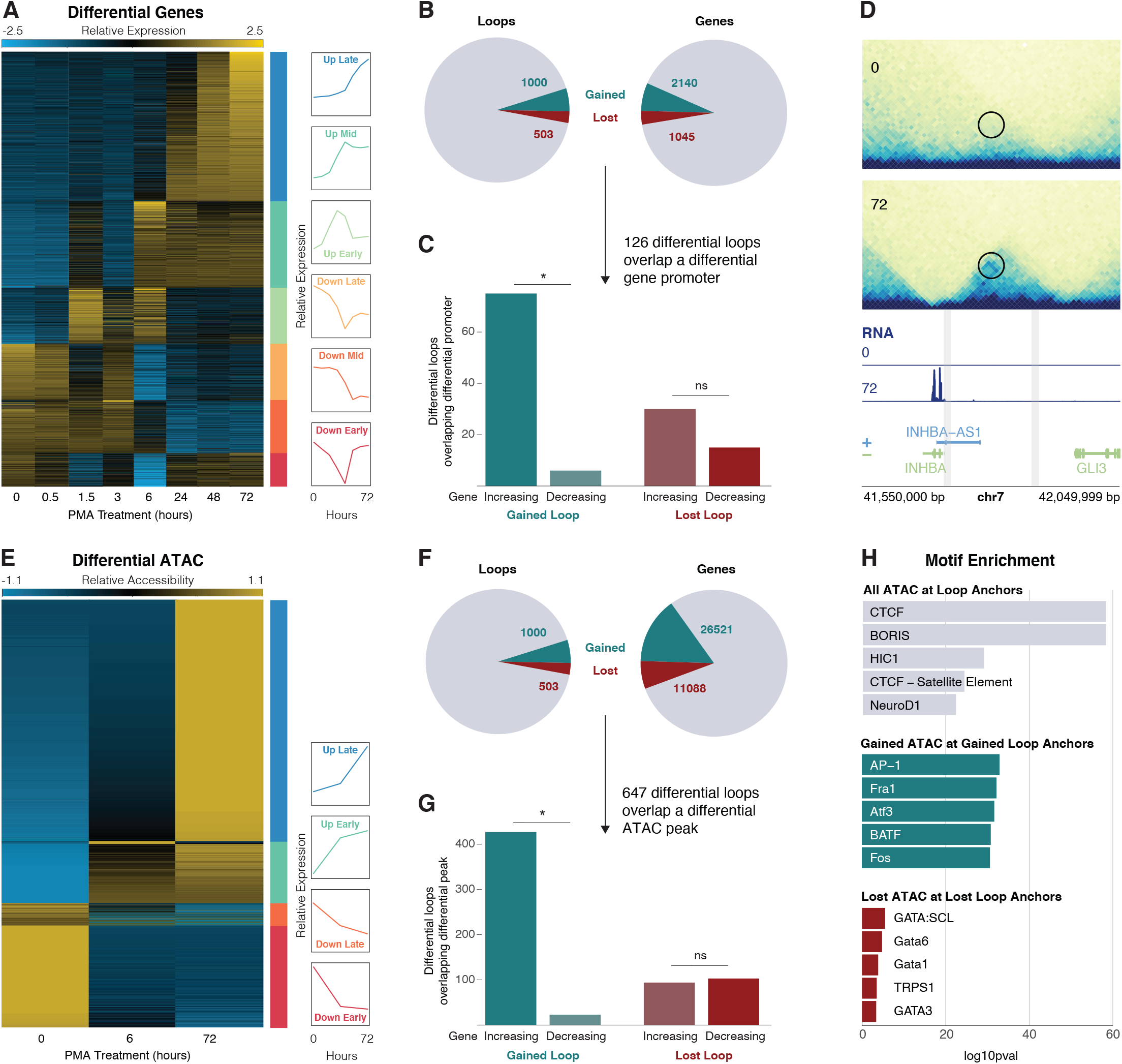
Gene expression and chromatin accessibility changes at differential loops. **(A)** RNA-seq normalized counts for all differential genes. Clusters are indicated by bars on the right side of the heatmap. Line plots show the mean expression per cluster. **(B)** Pie charts showing the proportion of differential loops (left) and genes (right). **(C)** Concordance analysis for the 126 differential loops that had a differential gene promoter at an anchor. Asterisks represent p < 0.05 (binomial test). **(D)** Example region of a gained loop with an increased gene at the INHBA locus. **(E)** ATAC-seq normalized counts for all differential peaks. Clusters indicated by bars on the right side of the heatmap. Line plots show the mean expression per cluster. **(F)** Pie charts showing the proportion of differential loops (left) and ATAC peaks (right). **(G)** Concordance analysis for the 647 differential loops that had a differential ATAC peak at a promoter. Asterisks represent p < 0.05 (binomial test). **(H)** TF motif enrichment analysis on all ATAC peaks at all loops (top), concordant gained ATAC peaks at gained loop anchors (middle), and concordant lost ATAC peaks at lost loop anchors (bottom).

To determine the relationship between differential loops and gene transcription, we intersected the promoters of differential genes with differential loop calls (**Fig 2B**). We found that 9% (126 of 1,503) of differential loop anchors overlapped a differential gene promoter. Interestingly, 93% (75 out of 81) of gained loops that overlapped a differential gene promoter showed the same direction of change as the gene (p = 2.91 × 10^−16^, binomial test) (**Fig 2C**). This is consistent with our previous work in macrophages^30,31^ and suggests that gained loops are relevant in increasing transcription of genes at their anchors. Upregulated genes found at the anchors of gained loops include *TGFB1* and *THBS1*, both of which are megakaryocyte-related^32,33^. An example of a gained loop with a concordant increase in gene expression is present at the *INHBA* locus (**Fig 2D**).

In contrast, lost loops that overlapped a differential gene promoter showed no such concordant behavior, with only 33% (15 out of 45) exhibiting the same directional change as the gene (p = 0.04, binomial test) (**Fig 2C**). These findings again agree with our previous work in macrophages^30,31^, where we did not see a significant decrease in expression of genes at lost loop anchors, suggesting that loss of looping is not sufficient to decrease transcriptional output^30,31^. This is also consistent with work by Rao et al that found almost no change in gene transcription following virtually complete abrogation of loop-extrusion^14^. Taken together, this suggests that loss of looping is generally not sufficient to induce a decrease in transcriptional output of genes at loop anchors. The anti-concordance we observed at loop anchors was also observed when looking at genes found between loop anchors. 45% of lost loops (228 out of 503) had a differential gene between their anchors, 64% of which (146 out of 228) overlapped a gene that was increasing (p = 2.69 × 10^−5^, binomial test) (**Fig S2C**). Genes that overlapped the interior of lost loops were expressed at significantly higher levels compared to genes within gained loops (p = 1.85 × 10-23, Wilcoxon rank sum test) (**Fig S2D**). This also agrees with our work in macrophages^30,31^ showing that extremely high expression of genes within loop boundaries was associated with a weakening of the loop, suggesting that high transcription at loop interiors might antagonize loop extrusion. Indeed, several other studies provide evidence that transcription can serve as a barrier to and/or interfere with loop extrusion^34–36^.

### Gained loops are associated with increased accessibility at AP-1 motifs

To assess which TFs were involved in loop based regulation, we mapped chromatin accessibility using ATAC-seq in K562 cells treated with PMA for 0, 6, and 72 h. Differential chromatin accessibility analysis with DESeq2 revealed 37,609 differential peaks (adjusted p-value, LFC > 2), which we grouped into 4 clusters based on whether the peak reached maximal or minimal normalized counts at 6 or 72h (**Fig 2E**). As was the case with gene expression, differential chromatin accessibility peaks were highly concordant at the anchors of gained, but not lost, loops (**Fig 2F-G**).

To understand the mechanisms driving differential looping we performed TF motif enrichment on various sets of ATAC peaks at loops anchors (**Fig 2H**). As expected CTCF was the most enriched motif at ATAC peaks overlapping all loop anchors, which is consistent with its known role in loop formation and maintenance. In contrast, peaks of gained chromatin accessibility at gained loop anchors were highly enriched for Activator Protein 1 (AP-1) family members. This is consistent with our previous work showing the enrichment of AP-1 at the anchors of gained loops during macrophage development^30^. Peaks of decreased chromatin accessibility at lost loop anchors were enriched for Gata family members, albeit to a far lesser degree compared to the enrichments observed at all or gained loops. GATA TFs have a well established role in mediating cell fate decisions in myeloid cell development^37–39^.

### Differential looping is associated with chromatin features both at and between loop anchors

To gain further insight into the mechanisms driving differential looping, we generated matched CUT&RUN datasets for multiple TFs and histone modifications. Due to the enrichment of AP-1 motifs at the anchors of gained loops, we performed CUT&RUN for JUN, a member of the AP-1 family. We also targeted histone H3 K27 acetylation, which is commonly used to identify putative enhancers, and CTCF and RAD21, proteins with known roles in DNA looping. For each assay, we performed differential analysis and compared how each differential feature corresponds to differential looping (**Fig S3A-D**). For all features, we observed statistically significant concordance at gained loop anchors (**Fig 3A-B**). With the notable exception of RNA, all features were also concordant at lost loop anchors, albeit to a lesser degree.

**Figure 3.**
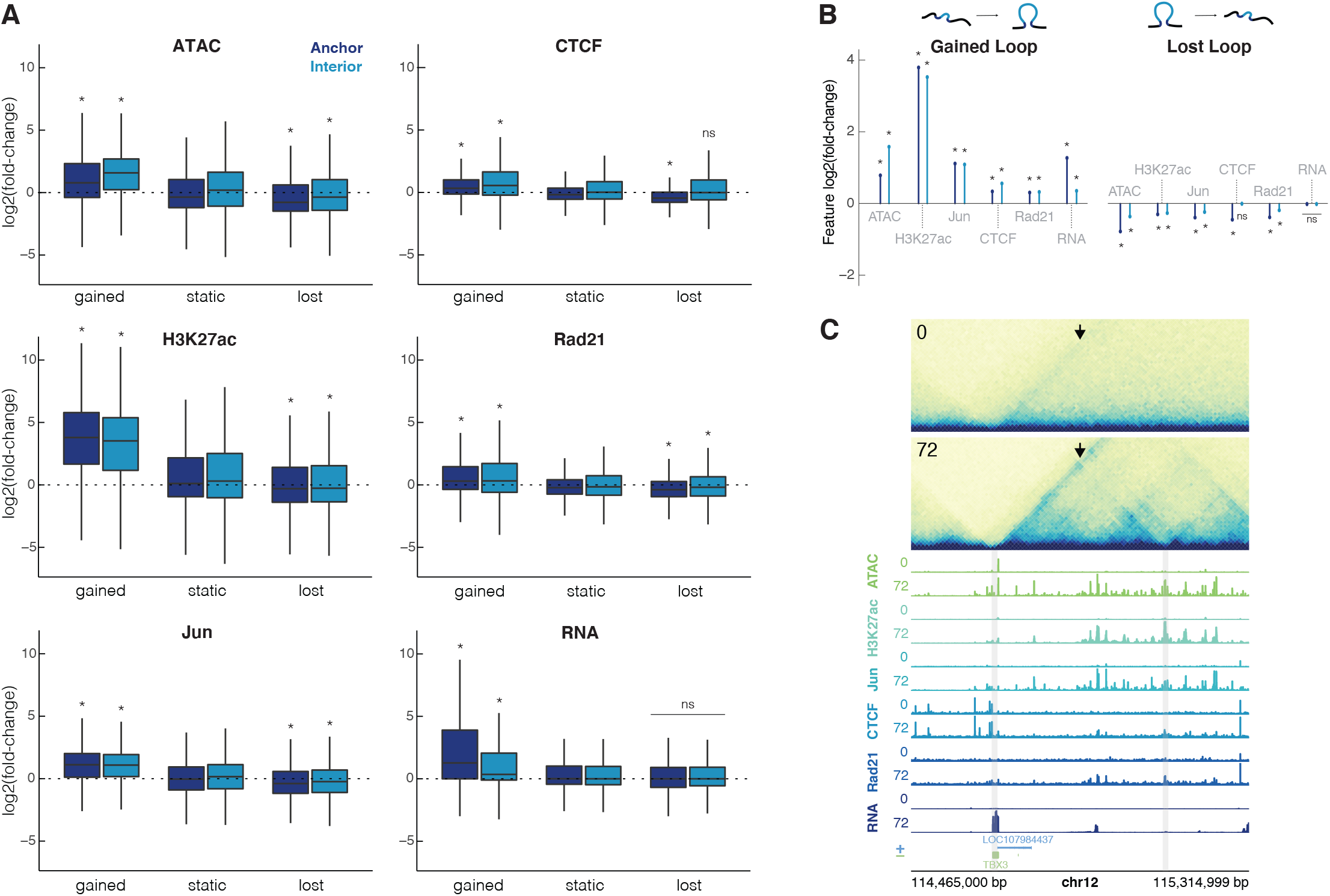
Changes in chromatin features are correlated with changes in looping. **(A)** Intersections of each feature at the lotop anchors and interior for gained, static, and lost loops (dark blue = anchor, light blue = interior). All plots are on the same scale for the y-axis, showing log2(fold-change). Wilcoxon rank sum test was performed for each feature to compare gained/lost anchors to static anchors and gained/lost interiors to static interiors, asterisks represent p < 0.05. **(B)** Median unshrunken log2(fold-change) of each dataset at gained loops (left), and lost loops (right). Asterisks represent p < 0.05, dots represent p > 0.05. **(C)** 600 kb region around a loop at the TBX3 locus at 10 kb resolution. The arrow is pointing to the gained loop. Signal tracks for ATAC-seq, H3K27ac, Jun, CTCF, Rad21, and RNA for K562s at 0h and differentiated megakaryocytes at 72h show increased occupancy for all features. Gray bars indicate 10 kb loop anchors.

Investigating these trends more closely revealed two prominent features. First, while all features increased at gained loop anchors, some exhibited bigger changes than others. Surprisingly, the canonical mediators of chromatin looping, CTCF and RAD21, increased by only 1.26- and 1.23-fold respectively at gained loop anchors. In contrast, histone H3 K27ac increased nearly 14-fold at gained loop anchors (median = 13.86-fold). Chromatin accessibility and JUN binding increased by modest amounts, 1.72- and 2.16-fold respectively. Second, while increased signals were observed at gained loop anchors, they were also observed between loop anchors. For example chromatin accessibility and histone H3 K27ac peaks within the boundaries of gained loops increased by 2.99- and 11.52-fold respectively.

Examples of these trends are evident at the *TBX3* locus (**Fig 3C**). Gained enhancer-promoter looping is associated with increases in CTCF and RAD21 occupancy at loop anchors as well as large gains in chromatin accessibility and JUN occupancy at, between, and even beyond loop boundaries. Expression of TBX3 itself, a TF known to regulate developmental transitions^40^, increases by over 50 fold. Taken together these results suggest that differential looping may involve more than just alterations to CTCF and RAD21 occupancy. They may also be mediated by chromatin modifying proteins and condition-specific TF binding events that act both at loop anchors and within the loop interior.

### Changes in chromatin features predict changes in chromatin looping

To explore this further and determine which chromatin features are the most predictive of changes in chromatin looping, we investigated how changes in each feature correlated with changes in looping (**Fig 4A**). Despite the relatively small fold change of CTCF peaks at differential loop anchors, we found that CTCF occupancy changes had the highest correlation (R^2^ = 0.189) with loop log2 fold-change (**Fig 4A**), consistent with the role of CTCF in the formation and maintenance of loops. However, changes to multiple features in the loop “interior”—the region in between the two anchors—also strongly exhibited strong correlations with differential looping. Interior chromatin accessibility, histone H3 K27 acetylation, and JUN occupancy had correlations of 0.175, 0.154, 0.127, followed by anchor RAD21 occupancy (R^2^ = 0.117). Several features exhibited very slight negative correlations including interior gene expression and CTCF occupancy, which is consistent with each of these having the ability to antagonize loop extrusions as previously described^13,34,41^. In addition to correlations with changes in looping, many of the features are highly correlated with each other (**Fig S4**).

**Figure 4.**
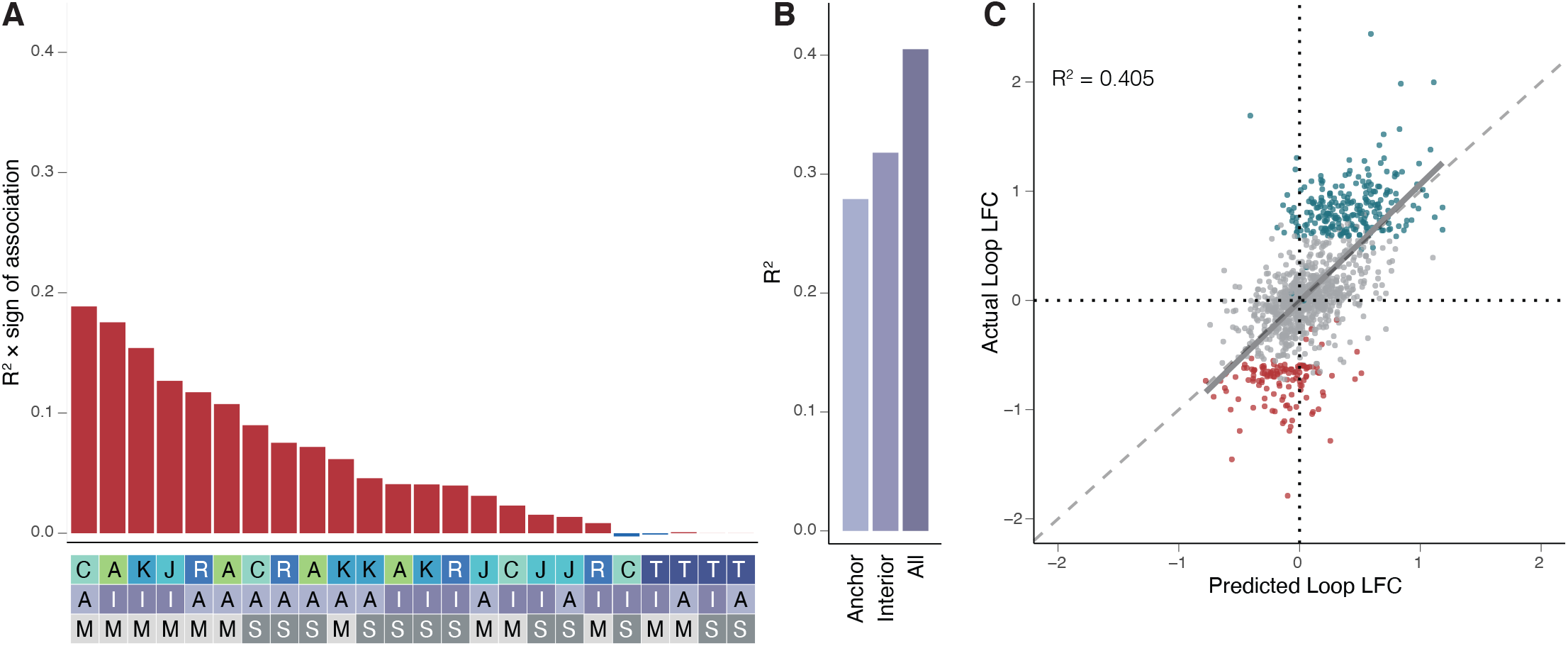
Changes in chromatin features predict changes in chromatin looping. **(A)** R^2^ multiplied by the sign of association for all possible features correlated individually against changes in all loops in the model (top). Heatmap (bottom) is a legend where the feature, position, and measure for each bar are reported. (features: A = ATAC, K = H3K27ac, J = JUN, C = CTCF, R = RAD21, T = RNA; position: A = anchor, I = interior; measure: M = max, S = sum). **(B)** R^2^ values calculated for the anchor only, interior only, or all feature models. **(C)** Scatterplot showing the predicted loop fold-change vs actual loop fold-change for the testing dataset for looping. (Gray = static loops, teal = gained loops, maroon = lost loops, R^2^ calculated for all loops included in the testing dataset).

Given the relationships between changes in looping and multiple features both at and between loop anchors, we next asked if we could build a better model to predict differential looping using multiple features simultaneously. We used the caret package to perform LASSO regression^42,43^. We selected 1,127 differential loops and 2,254 non-differential loops that were matched for distance and contact frequency as our training set and held out 376 differential loops and 752 non-differential loops as a testing set to evaluate our model. We then used LASSO with 10 cross-validations to generate a predictive model using various sets of features (**Fig 4A**) and evaluated it by applying it to our test set. Using all anchor features improved the correlation to R^2^ = 0.279, far higher than using the best single feature alone (CTCF anchor max, R^2^ = 0.189; **Fig 4B**). Surprisingly, using only interior features yielded an even more accurate model with an R^2^ of 0.317 (**Fig 4B**). Combining all features at both the anchors and interiors produced the strongest predictions with an R^2^ of 0.405 (**Fig 4B-C)**. Further, the model accurately predicted the signs of 91% of gained loops and 72% of lost loops. The predictive power of histone H3 K27 acetylation, chromatin accessibility, and JUN occupancy is surprising, and especially so given that the highest correlations were observed within loop boundaries rather than at anchors themselves. This may suggest that epigenetic changes between anchor sites play a significant role in modulating loop strength.

### Changes in and histone acetylation and chromatin structure predict changes in gene expression

Predicting gene expression patterns from chromatin features is a long standing and difficult problem in the field of gene regulation. Recent advances have been made by incorporating both 2D (e.g. histone H3 K27 acetylation) and 3D (e.g. Hi-C contacts) features into the Activity-By-Contact model of gene regulation^44^. This model has been successful in assigning enhancers to their target genes but has typically been applied to resting cells rather than biological transitions. A notable exception is Beagan et al. that correlated changes to ABC score to changes in gene expression^45^ albeit at only a handful of genomic loci. We leveraged this approach to determine if chromatin dynamics could help predict changes in gene expression in a genome-wide fashion.

To evaluate our ability to predict changes in gene expression we built and evaluated four linear models. In the first model, changes in gene expression were predicted based solely on changes in promoter H3 K27ac which performed well with an R^2^ of 0.401 (**Fig 5A**). In the second model, adding information from the nearest enhancer slightly decreased the predictive power (Wilcoxon rank sum test on permutations, p = 1.99 × 10^−6^; median R^2^ = 0.401; **Fig 5B**). In contrast, in the third model, by combining H3 K27ac information from both the promoter and enhancers that were physically looped to the promoter, the R^2^ increased significantly to 0.421 (Wilcoxon rank sum test on permutations, p = 1.99 × 10^−122^; **Fig 5B**). We then calculated a modified ABC score (see methods) by taking the product of loop strength and distal enhancer activity for each looped enhancer-promoter pair, summing across all enhancers that were looped to each gene, and then calculating the fold-change. Finally, the last linear model built using both promoter histone H3 K27ac and change in ABC score increased the R^2^ even further to 0.467 (**Fig 5D**), a significant improvement compared to all other models (Wilcoxon rank sum test on permutations, p = 1.95 × 10^−182^; **Fig 5E-F**). The improvement was even more drastic when building and applying the model to specifically differential genes and differential loops. For the models built on differential genes and loops, R^2^ values improve from 0.674 (promoter only model) to 0.763 for the promoter plus ABC score model (**Fig S5A-E**). These findings suggest that alterations to both enhancer activity and contact frequency can tune transcriptional programs during cellular differentiation.

**Figure 5.**
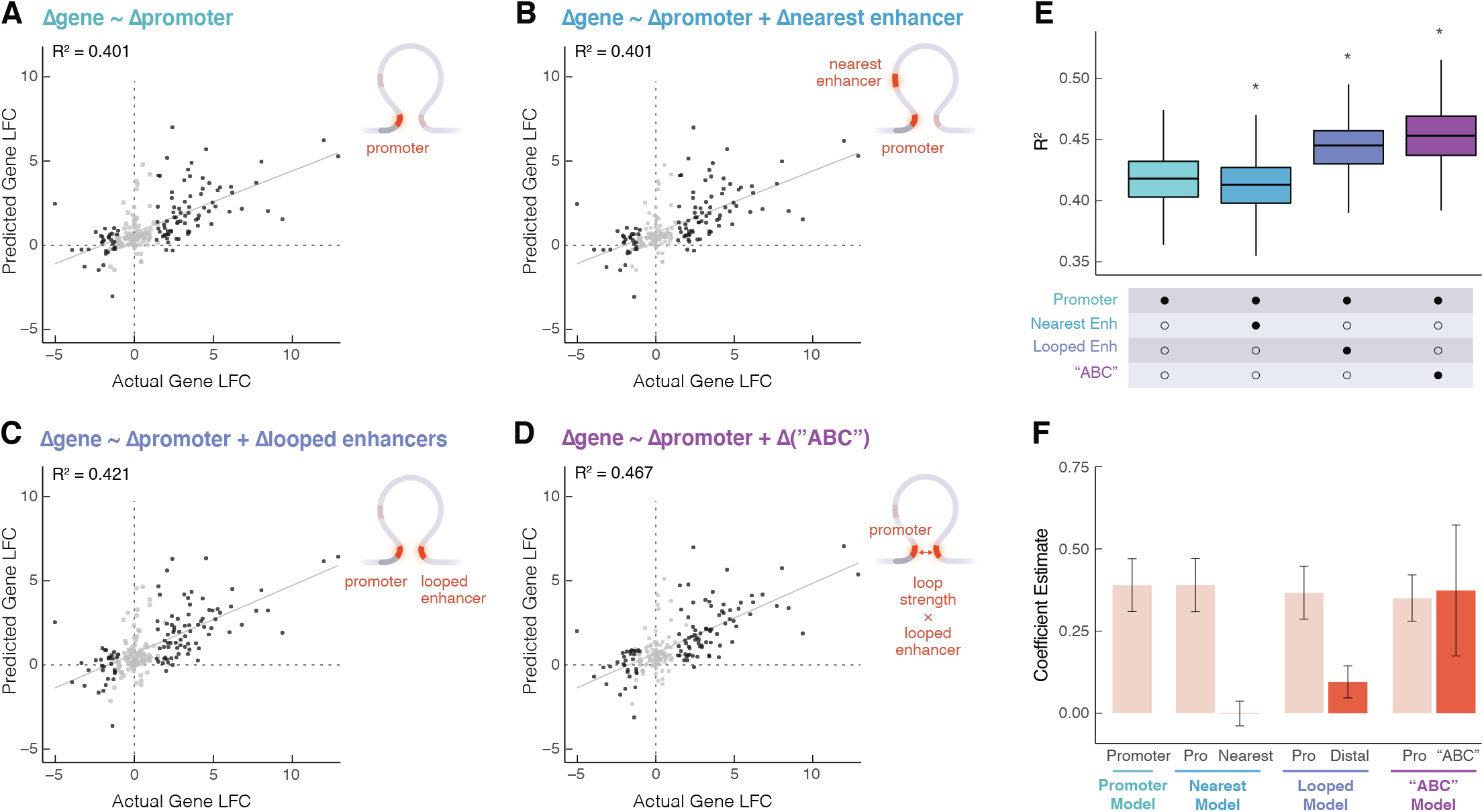
Changes in gene expression are explained by combined proximal and distal enhancer activity and loop strength. Scatter plots showing predicted gene-fold change vs actual gene fold-change based on **(A)** promoter H3K27 ac LFC alone, **(B)** promoter H3K27ac LFC and the nearest enhancer to the promoter’s LFC, **(C)** promoter H3K27ac LFC and distal looped H3K27ac, and **(D)** promoter H3K27ac LFC and the modified ABC LFC (dark gray = differential gene, light gray = static gene). **(E)** R^2^ for each model calculated based on 1000 permutations of splitting data into training and testing sets. Wilcoxon rank sum test was performed to compare each group to the promoter only model, asterisks represent p < 0.05. **(F)** Estimates for each term in each model were calculated based on 1000 permutations of splitting the data into training and testing sets.

## DISCUSSION

Collection and integration of deeply sequenced Hi-C and other genomic data characterizing the differentiation of K562 cells into a megakaryocyte-like state strengthened previous findings and provided novel observations into the mechanisms and impacts of changes to 3D chromatin structure. Our results confirmed canonical roles for CTCF and RAD21 in loop establishment and were consistent with chromatin looping playing a role in transcriptional activation and/or enhancement; however, these results also revealed strong correlations between differential looping and other regulatory features including chromatin accessibility, histone H3 K27 acetylation, and AP-1 occupancy and found a lack and/or anti-correlation of gene expression at lost loops. Intriguingly, differential looping correlated with transcriptional and regulatory features both at and between loop boundaries. In many cases, the correlations with internal features were stronger than the correlation with anchor signals. These results raise further questions regarding the mechanisms driving differential looping and point to previously underappreciated roles of molecules other than CTCF and cohesin.

The correlation between gained loops and the expression genes at their anchors agrees with many previous studies and supports the role of chromatin looping in gene activation. The increased predictive power when incorporating acetylation dynamics of looped enhancers and the dynamics of loop strength itself further emphasize this point. In contrast, loss of looping does not coincide with decreased expression of anchor genes. In fact, more genes were increasing in expression than decreasing at the anchors of lost loops. This is consistent with our previous studies of macrophage development and activation, neither of which identified a decrease in expression at lost loop anchors^30,31^. Taken together, this suggests that while loops may be involved in transcriptional activation, loss of looping alone may not be sufficient for transcriptional repression or decreased expression. This agrees well with previous studies involving rapid depletion of cohesin and subsequent global loss of loop extrusion-driven loops. Rao et al^14^ found that loop elimination did not substantially alter gene transcription—as measured directly using PRO-seq—in human colorectal cancer (HCT-116) cells. In contrast, loop disruption does inhibit activation of proinflammatory transcription in macrophages treated with LPS^46^ (a bacterial cell wall component commonly used as a model for inflammatory activation), again suggesting that loops do play a role in gene activation. While it is difficult to fully reconcile these findings, the evidence seems to be mounting that while increased looping can play a role in gene activation, events beyond loop disruption are required to decrease expression levels. One possible explanation is that looping imparts some sort of regulatory memory that is not erased as soon as the loop is disrupted. Similar mechanisms (e.g. the kiss-and-run mechanism)^47^ have been suggested before. While these trends are coming into focus, more functional studies are required to understand the exact role of chromatin looping in gene activation and repression.

Our data also support the theory that loop loss may be a result, rather than cause, of changes in gene transcription. Differential genes within lost loops are significantly biased towards increased expression. And those increased genes tend to be fairly highly expressed. We observed this exact same phenomena in macrophages responding to LPS^31^. This is consistent with transcription occurring at a high level between loop boundaries being antagonistic to loop extrusion, something that is easy to imagine given that both involve fairly large molecular complexes traversing the same stretch of DNA. Indeed several previous studies have suggested that transcription can act as a molecular barrier to the loop extrusion process^34–36^.

Interestingly, we revealed that changes to multiple other chromatin features (i.e. chromatin accessibility, histone H3 K27 acetlyation, and AP-1 occupancy) are roughly as predictive as changes to known loop extrusion-related proteins CTCF and RAD21. The correlation between looping and these other features was strongest for features within the loop interior which may provide clues into the nature of this relationship. One intriguing possibility is that increased accessibility, histone H3 K27 acetylation, and AP-1 binding might play a role in increasing the efficiency or rate of cohesin loading. This seems consistent with previous findings that suggested that cohesin loading takes place preferentially at active gene promoters^4^ which are also associated with histone acetylation, TF binding, and chromatin accessibility. It is important to acknowledge that we can not rule out the possibility that all of these internal changes are the result, rather than cause, of differential looping events.

Despite the intriguing findings presented here, several limitations of this study must be considered when interpreting the data and speculating about their meaning. First, this study is largely correlative. While intersecting the results of this study can provide important mechanistic insights, several of the results raise new questions that must be addressed by future functional experiments. Second, in contrast to our previous work^31^, this time-course lacked the temporal resolution to put regulatory and transcriptional events in temporal order, which makes it difficult to infer the direction of causality between any two features. This was a conscious decision as the current cost of sequencing makes it unfeasible to acquire deeply sequenced data sets across deeply sampled time courses; however, this is likely to change soon as sequencing costs continue to decrease^48,49^. Finally, while this study encompassed a broad number of regulatory features—including ATAC-seq, which when combined with motif analysis can provide insights into the binding patterns of hundreds of TFs—there are a vast number of other features that likely influence looping (e.g. DNA methylation) for which we are not measuring nor explicitly accounting for.

Despite these limitations, these findings improve our understanding of how different trans regulators and epigenetic features govern changes in looping, as well as our understanding of the relationship between looping and gene expression. The deeply sequenced nature of this differential Hi-C analysis offers a uniquely well-powered dataset with which to explore a pressing number of biological questions. Moverover, this data was acquired in one of the most widely studied human cell lines (K562) for which hundreds of publicly available genome-wide data sets are already available. As such, this study provides a valuable new resource for future studies of chromatin biology.

## DATA AVAILABILITY

Raw and processed data for Hi-C (GSE213909), RNA-seq (GSE213386), ATAC-seq (GSE213295), and CUT&RUN (GSE213908) are publicly available on GEO and SRA under SuperSeries 213909.

## ACKNOWLEDGMENTS

We thank Erika Deoudes for graphic design of figures and preparation of the manuscript. We thank Sam Pattenden for allowing us to use the Covaris LE220 instrument. We thank Katie Reed for assistance with project design and intellectual discussions. We thank Nicole Kramer for the development of plotgardener and help with generating figures and data visualization.

## AUTHOR CONTRIBUTIONS

M.L.B. designed and performed most of the experiments, performed all of the computational analysis, and wrote the manuscript.

E.S.D. developed some of the software used and assisted with computational analysis.

I.Y.Q. prepared ATAC-seq libraries.

M.I.L. oversaw data analysis and linear modeling

H.W. acquired funding, supervised data analysis, and helped write the manuscript.

D.H.P. acquired funding, conceptualized the project, supervised experiments and data analysis, and helped write the manuscript.

## METHODS

### K562 culture and differentiation

K562 were cultured in RPMI media (Corning, cat # 10-040-CV) with 10% fetal bovine serum (FBS) (Gibco, cat # 26140079) and 1% penicillin-streptomycin (PS) (Gibco, cat # 15140122). For megakaryocyte differentiation, K562 were plated in either 6-well plates (RNA-seq, ATAC-seq) or T-175 flasks (Hi-C, CUT&RUN) at a density of 1 × 10^5^ cells/mL and treated with 25 nM PMA (Sigma-Aldrich, cat # P1585-1MG). After 24h, the cells become semi-adherent. Cells were provided with fresh media and PMA after 24h and 48h. Cells were collected without treatment or after 6 or 72h. For all treatments and library preparations, K562s were thawed and immediately split into two T-25 flasks to create biological replicates.

### Crosslinking

Cells were cultured in T-175 flasks containing 10 × 10^6^ cells for Hi-C and 5 × 10^6^ cells for CUT&RUN at 1 × 10^5^ cells/mL. The collection protocol differs for where the cells are at during the differentiation. <u>For suspended cells (0h and 6h)</u>: Media was centrifuged at 300 × g for 5 min. Cells are in suspension for 0h and 6h. Suspended cells were collected and spun down 300 × g for 5 min. Pellets were resuspended in 10 mL of 1% formaldehyde (Thermo Fisher Scientific, cat # 28908) in RPMI for 10 min with rotation. <u>For semi-adherent cells (72h)</u>: The suspended cells were harvested and centrifuged for 300 × g for 5 min. Pelleted cells were resuspended in 10 mL of 1% formaldehyde, and added to the adherent cells in the T-175 flask to crosslink all cells at once, where they were put on the shaker for 10 min (from here, the protocol is the same for all time points). Cells were quenched with cold glycine for 5 min (Invitrogen, cat # 15527013) to a final concentration of 2.0 M for 5 min. Cells were then centrifuged at 562 × g, resuspended in cold PBS (Corning, cat #21040CV), and split into 3 tubes of approximately 3 × 10^6^ cells each (HiC) or 10 tubes of approximately 5 × 10^5^ each (CUT&RUN). Cells were spun again at 562 × g for 5 min and washed again with cold PBS, then aspirated and flash frozen in liquid nitrogen, and stored at −80°C.

### In situ Hi-C library preparation

Four treatments (biological replicates) were performed. For each treatment, either two or three frozen pellets (3 × 10^6^ each) were used to generate technical replicates (3 technical replicates for the first two biological replicates and 2 technical replicates for the last two biological replicates). Libraries were generated according to the protocol as described in Rao et al^15^. Briefly, crosslinked we lysed the cells, isolated nuclei, and used MboI (New England Biolabs, cat # R0147L) to digest chromatin overnight. The fragment ends were biotinylated, proximity ligated, and reverse crosslinked. Quantification of shearing DNA was achieved with Qubit (dsDNA Broad Range (BR) assay) (Thermo Fisher Scientific, cat # Q32850). We then sheared the samples on a Covaris LE 220 (duty factor 25, PIP 500, 200 cycles/burst, 90 seconds). 2% of each sample was run on a 2% agarose gel to confirm fragmentation. Size selection with AMPure XP beads (Beckman Coulter, cat # A63881) was then performed to select for DNA fragments between 300 and 500 bp. Biotinylated chromatin was pulled down with streptavidin beads. Biotin was then removed from unligated ends and the libraries were end repaired. We added the Illumina TruSeq Nano (Set A) (Illumina, cat # 20015960) indices to each sample in a combination appropriate for pooling and amplified using 9 cycles of PCR. Final quantification was achieved using Qubit (dsDNA High Sensitivity (HS) assay) (Thermo Fisher Scientific, cat # Q32851) and Tapestation (D1000 screentape) (Agilent, cat # 5067-5584). Libraries were pooled to 10 nM and sequenced across 7 Illumina NovaSeq S4 lanes (Novogene, 150-bp paired-end).

### RT-qPCR

We extracted RNA from 5 × 10^5^ cells using the QIAGEN RNeasy Mini kit (Qiagen, cat # 74014) with DNase I treatment (Qiagen, cat # 79254) and quantified with a Qubit Broad Range assay (Thermo Fisher Scientific, cat # Q32850). Reverse transcription into cDNA was performed with the iScript cDNA synthesis kit (Bio-Rad, cat # 1708891). qPCR was performed with the TaqMan reagents using probes for ITGB3, KLF1, and GAPDH (Thermo Fisher Scientific, cat # Hs01001469, Hs00610592, Hs02786624).

### RNA-seq library preparation

We extracted RNA from 5 × 10^5^ cells using the QIAGEN RNeasy Mini kit with DNase I treatment. To confirm quality of libraries, we checked RNA integrity numbers with a Tapestation RNA screentape (Agilent, cat # 5067-5577) and confirmed them all to be above 9.7. We determined the concentration of all RNA samples with the Qubit Broad Range assay (Thermo Fisher Scientific, cat # Q10211).

The KAPA RNA HyperPrep kit with RiboErase (HMR) (Kapa Biosciences, cat # KK8560) was used for library preparation. Illumina TruSeq adapters (Illumina, cat # 20015960) were diluted and 0.0075 nmol was added to each sample. We determined library concentration and fragments size with Qubit (dsDNA HS assay) and Tapestation (D1000 screentape). Libraries from each timepoint were pooled at 10 nM and each biological replicate was sequenced on an Illumina NextSeq 500 (75-bp paired-end, high output kit).

### ATAC-seq library preparation

We used the Omni ATAC-seq protocol as described in Corces et al^50^ with some adjustments to perform ATAC-seq. Two treatments (biological replicates) were performed. For untreated and 6h, cells were harvested and centrifuged at 500 × g for 5 min. For semi-adherent 72h cells, all of the floating cells were harvested. Adherent cells in each well were washed with 2 mL PBS, lifted with 500 μL 0.5 M EDTA for 5 min, and quenched with 3 mL of RPMI before combining with the floating cells. 5 × 10^5^ cells were used for library preparation. Illumina Nextera XT indices (Illumina, cat # FC-131-1001) (3.75 μL/sample) were used for PCR.

After 5 PCR cycles, 5% of each sample was used in qPCR to determine how many more cycles were necessary. We found that 4-7 cycles were sufficient for final amplification. AMPure XP beads were used to perform a final cleanup (0.5X followed immediately by 1.3X) and quantified with the Qubit (ds DNA HS assay). The concentration in molarity of samples was determined by the KAPA Library Quantification Kit (Kapa Biosystems, cat # 4854). Each replicate was pooled to 4 nM and sequenced separately on an Illumina NextSeq 500 (75-bp paired-end, high output kit).

### CUT&RUN library preparation

We generated CUT&RUN libraries following existing protocols^51^, but modified for the use of crosslinked cells. Cells were centrifuged at 500 × g at 4°C for 10 min. For H3 K27ac, 0.5 μL of 1:10 diluted antibody (Abcam, cat # ab4729) was added to each sample. For CTCF, 0.5μL of 1:10 diluted antibody (Thermo Fisher Scientific, cat # MA5-31344) was added to each sample. For JUN, 2.08 μL of stock antibody (Thermo Fisher Scientific, cat # MA5-15172) was added to each sample. For RAD21, 0.625 μL of stock antibody (Abcam, cat # ab992) was added to each sample. We then added 5 μL of KAPA Unique Dual-Indexed Adapters (Roche, cat # 08861919702) diluted to 750 nM. Libraries from each timepoint for H3 K27ac, CTCF, and JUN were pooled to 6 nM, and sequenced on an Illumina NextSeq 500 (75-bp paired-end, high output kit). RAD21 libraries from each timepoint were pooled to 9 nM, and were sequenced on an Illumina NextSeq 500 (75-bp paired-end, high output kit).

### Hi-C data processing and calling compartments, domains, and loops

We processed our Hi-C data using a modified version of the Juicer pipeline (version 1.9.8)^15^. Hi-C contact maps were generated at 5, 10, 25, 50, 100, 200, 250, 500, 1000, and 2500 kb resolution for each individual technical replicate that was sequenced. This was for 4 biological replicates, 3 timepoints, and 2-3 technical replicates each, totaling 30 unique samples. Additionally, all of the samples for each timepoint were merged to create merged Hi-C maps. All samples and replicates across all timepoints were also merged to create a “Mega” map.

Compartments were identified using the EigenVector R package at a 10 kb resolution^25^.

TADs were identified using the arrowhead command within the Juicer pipeline at 25 kb resolution. Cell type specific TADs were identified by merging with the mariner R package and using the denovo function.

Loops were called from the merged timepoint Hi-C files and Mega map with SIP^26^ (version 1.6.1). The settings “-g 2 −5 2000 -fdr 0.05” were used both on the timepoint and the Mega map. Loops were merged in R with mariner using the mergeBedpe function, providing a list of 33,914 loops.

A count matrix was prepared using mariner^52^, where unnormalized counts at each loop pixel from each technical replicate were extracted.

The compartment, TAD, and loop-level Hi-C maps were SCALE normalized and visualized with plotgardener^53^ at 100, 10, and 5kb resolutions respectively.

### Differential loop and aggregate peak analysis

DESeq2 was used to identify differential loops using the count matrix prepared as described above. Loops with a median count of 5 counts or less were filtered out. Counts from the technical replicates from each biological replicate were summed together and the design “~rep + time” was used, with a reduced design of “~rep” used to form a likelihood ratio test (LRT). Apeglm was used to calculate log-2 fold changes for each loop, comparing both 6 and 72h to 0h. Loops were deemed significant if they had an adjusted p-value < 0.05 and a log-2 fold change > 1.5.

Aggregate peak analysis (APA) was performed with mariner. For all, gained, and lost loops, the loop pixel and 10 pixels around the loop were extracted with SCALE normalization at 10 kb resolution.

### Hi-C Power Analysis

Power analysis was performed with the RNAPower package^27^. Dispersion was calculated from the differential loop analysis in DESeq2 as described above, where the minimum dispersion value was used. Power was modeled across various theoretical sequencing depths and replicates for identifying a log-2 fold change of 2 with a p-value of 0.05. The rnapower function was used with an alpha of 0.05/33914 and a cv of the square root of the dispersion value.

We subsampled our Hi-C data from the merged_nodups files to approximate sequencing depths of 100, 300, 500, and 700M per biological replicate. We then repeated our differential loop analysis using the subsampled data for either 2, 3, or 4 replicates using all of the same parameters and loop calls.

### RNA-seq processing

Fastq quality was assessed using the FastQC and MultiQC tools (FastQC version 0.11.5, MultiQC version 1.5)^54,55^. Fastq files were trimmed with Trim Galore! (version 0.4.3) and quantified with Salmon (version 1.4.0) to the hg38 genome^56,57^. Alignment was performed using HISAT2 (version 2.1.0), which generated BAM files that were indexed using samtools (version 1.9)^58,59^. The BAM files for each timepoint’s two biological replicates were merged using samtools and converted to bigwigs using deeptools (version 3.0.1)^60^ for easy visualization of signal tracks. Reads were summarized into a format compatible with DESeq2 using txImport (r version 3.3.1, tximport version 1.2.0)^29,61^.

### Differential gene analysis

DESeq2 was again used to identify differential genes. The txi file was used as input, and the DESeqDataSetFromTximport was used with “~rep + time” as the design. A reduced design of “~rep” was used to form an LRT, as previously for peak analysis. Shrunken log-2 fold change values were calculated for each gene by comparing the counts at each time point to 0h with apeglm^62^. Significant genes had an adjusted p-value < 0.05 and a log-2 fold change > 2.

The DESeq2 dataset was normalized with variance stabilized transformation. We then filtered for differential genes and calculated Z-scores based on standard deviation and mean. Replicates were then averaged, and kmeans clustering was used to identify 6 temporal clustered based on the vectors of Z-scores.

### ATAC-seq processing and peak calling

Adapters were trimmed and low quality reads were filtered out using Trim Galore! (version 0.4.3)^56^. BWA mem (version 0.7.17) was used to align reads to the hg38 genome and sorted using Samtools (version 1.9)^59^. PicardTools (version 2.10.3) was used to remove duplicate reads^63^. Mitochondrial reads were filtered out with Samtools idxstats^59^. For each timepoint, biological replicates were merged and indexed with Samtools. We called peaks were called on the merged files using MACS2 with the following parameters: -f BAM -q 0.01 -g hs --nomodel --shift 100 --extsize 200 --keep-dup all -B --SPMR (version 2.1.1.20160309)^64^. A comprehensive peak list was generated by merging peaks across all time points (181,136 peaks). For each peak across all biological replicates independently, counts were extracted with bedtools multicov, which was the input for differential peak analysis^65^. Signal tracks were generated from merged time points with deeptools (version 3.0.1) for visualization^60^.

### CUT&RUN processing peak calling

Adapters were trimmed off and low quality reads were filtered out with Trim Galore! (Version 0.4.3)^56^. BWA mem (version 0.7.17) was used to align reads to the hg38 genome, sorted with Samtools (version 1.9), and filtered for duplicates with PicardTools (version 2.10.3)^59,63^. Samtools was again used to index all BAM files. For each timepoint, biological replicates were merged and indexed with Samtools. We called peaks on the merged files using MACS2 with the following parameters: -f BAM -q 0.01 -g hs --nomodel --shift 0 --extsize 200 --keep-dup all -B --SPMR (version 2.1.1.20160309)^64^. A comprehensive peak list was generated for each antibody by merging peaks across all time points (X, Y, Z peaks for X, Y, Z datasets). For each biological replicate independently, counts were extracted with bedtools multicov, which was the input for differential peak analysis^65^. Signal tracks were generated from merged time points with deeptools (version 3.0.1) for visualization^60^.

### Differential ATAC-seq and CUT&RUN Peak Analysis

DESeq2 was again used to identify differential peaks from ATAC-seq and all CUT&RUN data. The count matrix generated in the ATAC-seq and CUT&RUN processing was used as the input with the function DESeqDataSetFromMatrix. We used “~rep + time” as the design and a reduced design of “rep” to form an LRT. Apeglm was used to calculate shrunken log-2 fold-changes for each peak for each dataset^62^. Significant peaks had an adjusted p-value of < 0.05 and an absolute log-2 fold change > 2. Peaks were clustered into up early, up mid, up late, down early, down mid, and down late by determining whether their max/min value was at 0, 6, or 72h.

### Gene Ontology and KEGG Pathway Enrichment Analysis

We used the HOMER function findMotifs.pl on each of our 6 gene clusters to identify enriched gene ontology terms and KEGG pathways with default settings^66^. For GO Terms, the biological_process.txt file was used and for KEGG pathways, the kegg.txt file was used.

### Motif Enrichment Analysis

We used the HOMER function findMotifsGenome.pl for all motif enrichment^66^. For motifs at the anchors of all loops, we intersected all ATAC peaks with all loop anchors. For motifs at the anchors of gained loops, we intersected all differential gained ATAC peaks with gained loop anchors. For motifs at the anchors of lost loops, we intersected all differential lost ATAC peaks with lost loop anchors. For each motif enrichment, all ATAC peaks were used as the background. The default parameters were used with the following adjustments: -size given.

### Genomic Intersections

The GenomicRanges and InteractionSet R packages were used to perform all genomic intersections^67^. Loop bedpe files were converted into GInteractions objects and were intersected with the coordinates for ATAC, H3 K27ac, JUN, CTCF, RAD21 peaks and genes with subset-ByOverlaps. The unshrunken fold-changes as calculated by DESeq2 analysis were extracted from each of the peaks that overlapped a loop anchor.

### Chromatin Looping Linear Model

The count matrices from previous analysis were again used to generate the data for the linear model. We used loop counts, peak counts from ATAC-seq and all CUT&RUN data, and transcripts per million (TPM) per 10 kb bin from RNA-seq. For all loops, each of these counts was extracted from both the anchors and each anchor individually. In the case of multiple peaks intersecting with loops, the sum of all peaks was recorded. We also recorded the maximum values from extracting counts from bigwig files instead. A final anchor measure was calculated by taking the product of signal (either sum or max) at both of the anchors for each loop. All interior measures were normalized to the length of the loop.

We added a pseudocount of 1000 to the entire data-frame (which is roughly 0.04 times the average count value) and then calculated the log(fold-change) between 72h and 0h. This “delta” matrix was then scaled, and DESeq2 log-2 fold-changes were used for looping. Our model consisted of all differential loops and twice as many static loops matched for distance and contact. Matched static loops were generated from the matchRanges^68^ function within the nullranges Bioconductor package. 75% of the dataset was used for training and the remaining 25% was reserved for testing.

Each feature was tested against loop LFC with the base R function lm to determine R^2^ values. The sign of correlation was determined with the cor function. LASSO regression was used to find a sparse model combining features, calling glmnet^42^ within the caret R package^43^. We trained the LASSO model on the training set, using anchor features only. We evaluated selected LASSO models on the test set using R^2^. This was repeated again for all interior features, and with all anchor and interior features combined.The R^2^ was calculated with the cor function. This was repeated for all interior features, and again repeated with all anchor and interior features combined.

### Gene Expression Linear Model

We used the lm function within the stats R package to model how gene expression changes correlate with changes in changes in proximal and distal acetylation, and looping. We used the GenomicRanges function subsetByOverlaps and linkOverlaps to determine which differential genes had promoter H3 K27ac and were looped to a distal H3 K27ac peak^67^, identifying 332 genes, and included 332 static genes matched for expression. We also identified the nearest enhancer with the nearest function of GenomicRanges. For proximal and distal H3 K27ac, we extracted the counts and calculated log2(-fold-change). For genes that had multiple enhancers, we took the sum of the counts at 0h and 72h and then calculated log2(fold-change). For the ABC score, we scaled all enhancer and loop counts to be between 1 and 100 to ensure that both factors were contributing equally to the interaction despite differences in sequencing depth. For each enhancer-promoter pair, we multiplied the normalized distal enhancer counts by the normalized loop strength counts, then calculated log2(fold-change). For genes with multiple enhancer-promoter pairs, we summed the multiplied score at 0h and 72h and then calculated log2(fold-change).

The first model only used changes in promoter acetylation to predict changes in gene expression. The following three models used promoter acetylation in addition to either nearest enhancer, distal enhancer, or the interaction between distal enhancers and loops. We trained on 60% of the data and tested on the remaining 40%. The predict function was used to predict on the testing dataset from the trained model and cor was used to calculate R^2^ values. For R^2^ and coefficient estimate calculations, we performed 1000 permutations of splitting the data into testing and training datasets.

## SUPPLEMENTAL FIGURES

**Figure S1.**
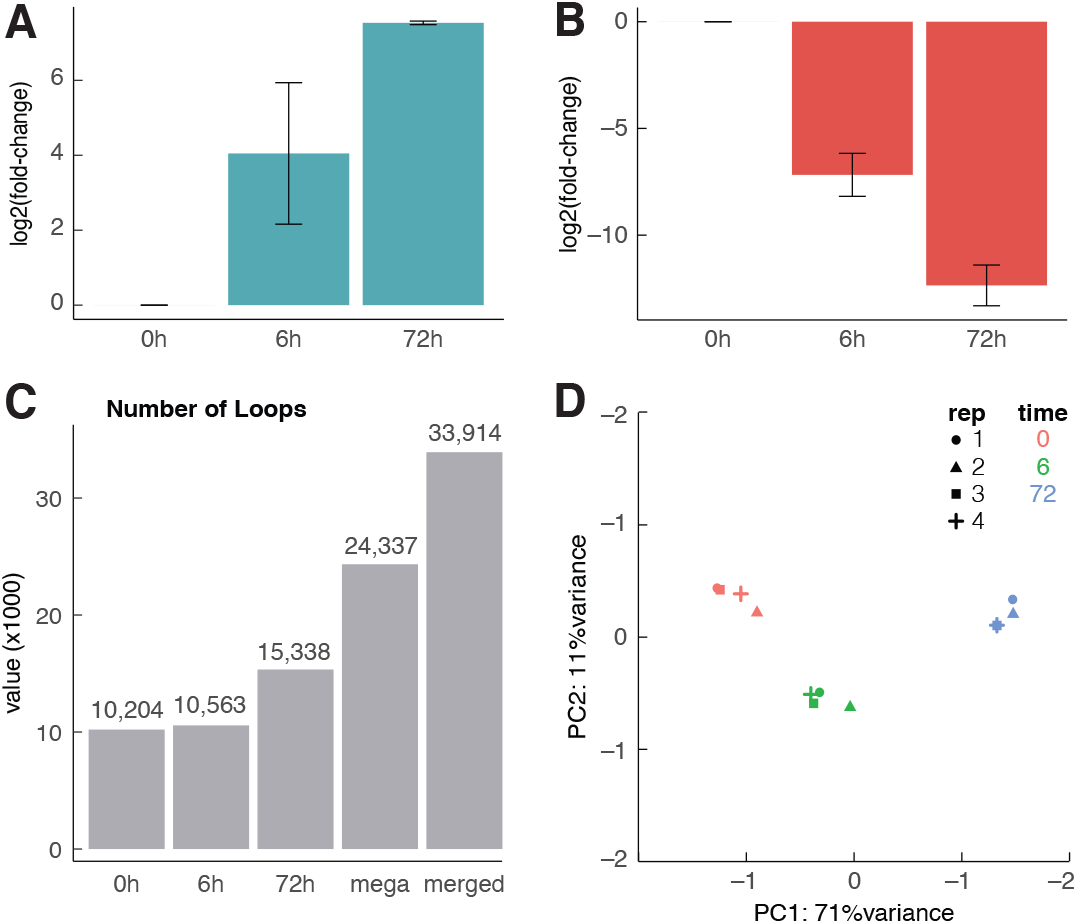
Confirmation of megakaryocyte differentiation and detection of loops. qPCR analysis of **(A)** ITGB3 and **(B)** KLF1 over megakaryocyte differentiation at 0, 6, and 72h. Two biological replicates and 4 technical replicates were collected, normalized to GAPDH levels, and log2(fold-change) was calculated relative to 0h. **(C)** Number of loops identified with SIP after 0, 6, or 72h of differentiation, after merging all timepoints together into the Mega map, and merging all timepoints together with mariner. **(D)** PCA plot showing similarities in loop counts between replicates and timepoints.

**Figure S2.**
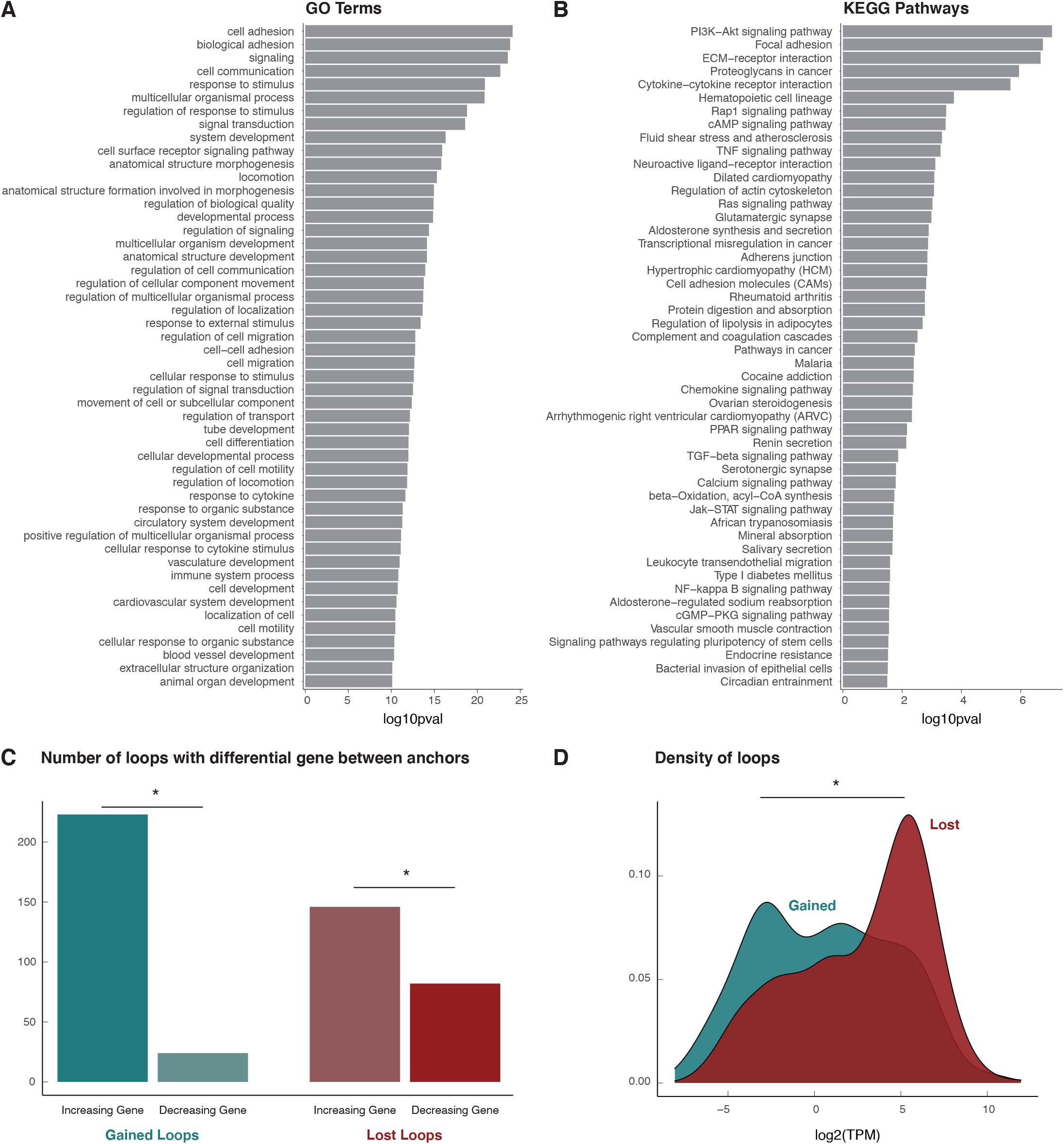
Megakaryocyte pathway enrichment and interior gene expression. **(A)** Top 50 GO terms for upregulated genes from RNA-seq. **(B)** Top 50 KEGG Pathways for upregulated genes from RNA-seq. **(C)** Concordance analysis for the 475 differential loops that had a differential gene promoter between their anchors. Binomial test performed for each comparison, *asterisks represent p < 0.05. **(D)** Expression of genes located between the anchors of gained and lost loops, *asterisk represents p < 0.05. (TPM: transcripts per million).

**Figure S3.**
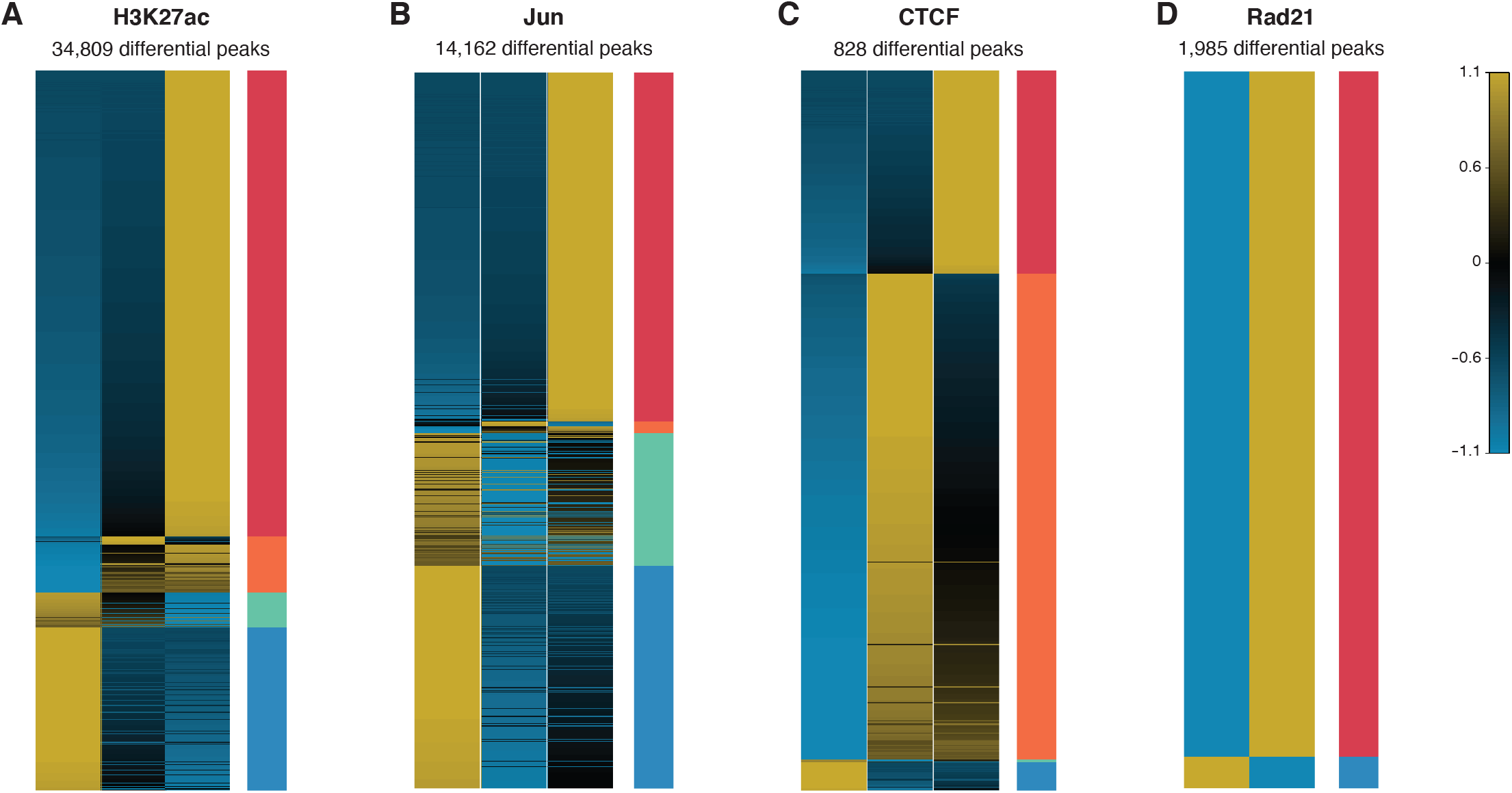
Differential chromatin and transcription factor binding events. Heatmaps showing normalized counts for **(A)** H3K27ac, **(B)** Jun, **(C)** CTCF, and **(D)** Rad21. Clusters are indicated by the bars on the right side of each heatmap, p < 0.05, log2(fold-change) > 2.

**Figure S4.**
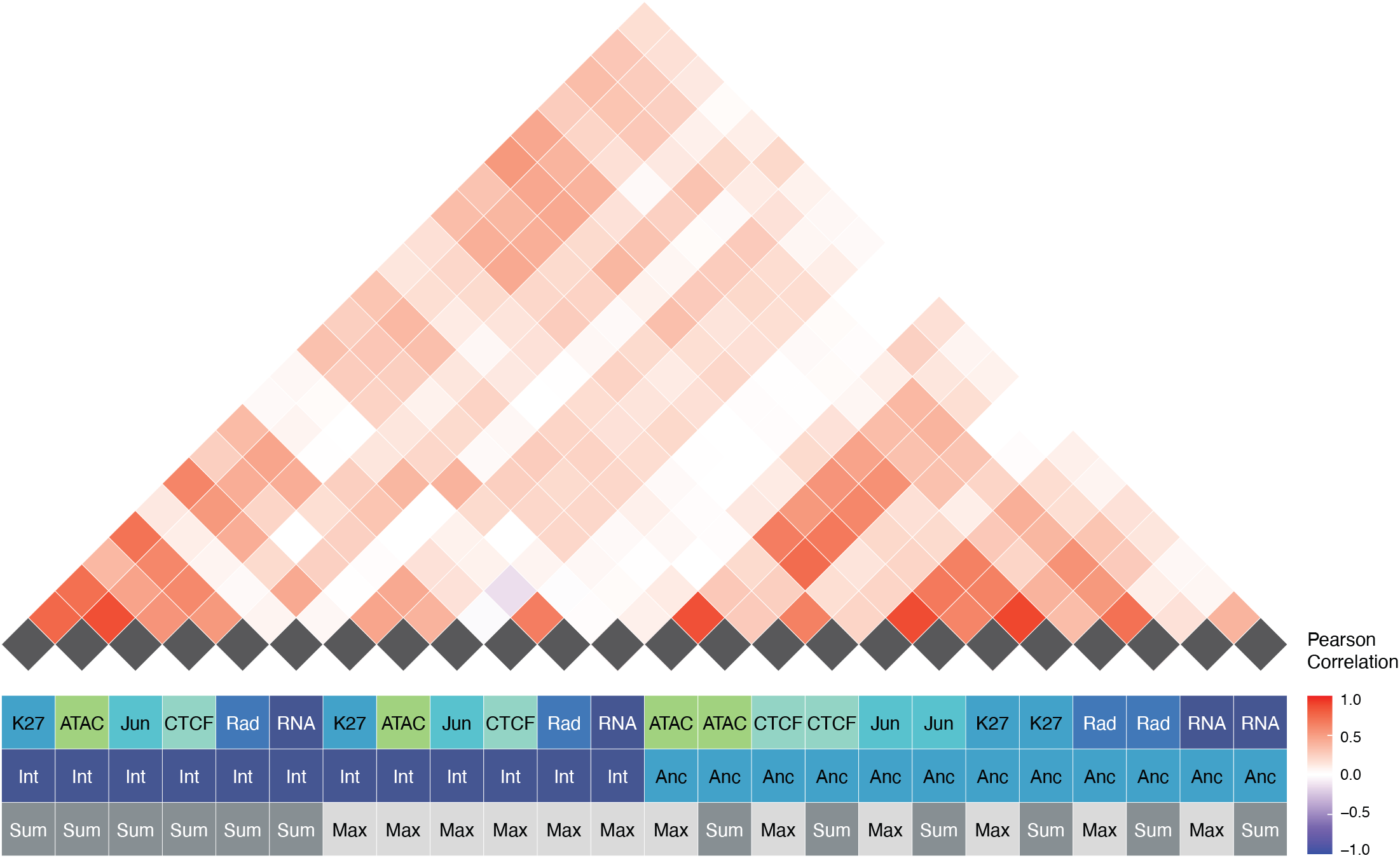
Correlation of genomic features. Correlation heatmap showing the individual correlations of each of the features (top) and legend representing which feature is represented (bottom).

**Figure S5.**
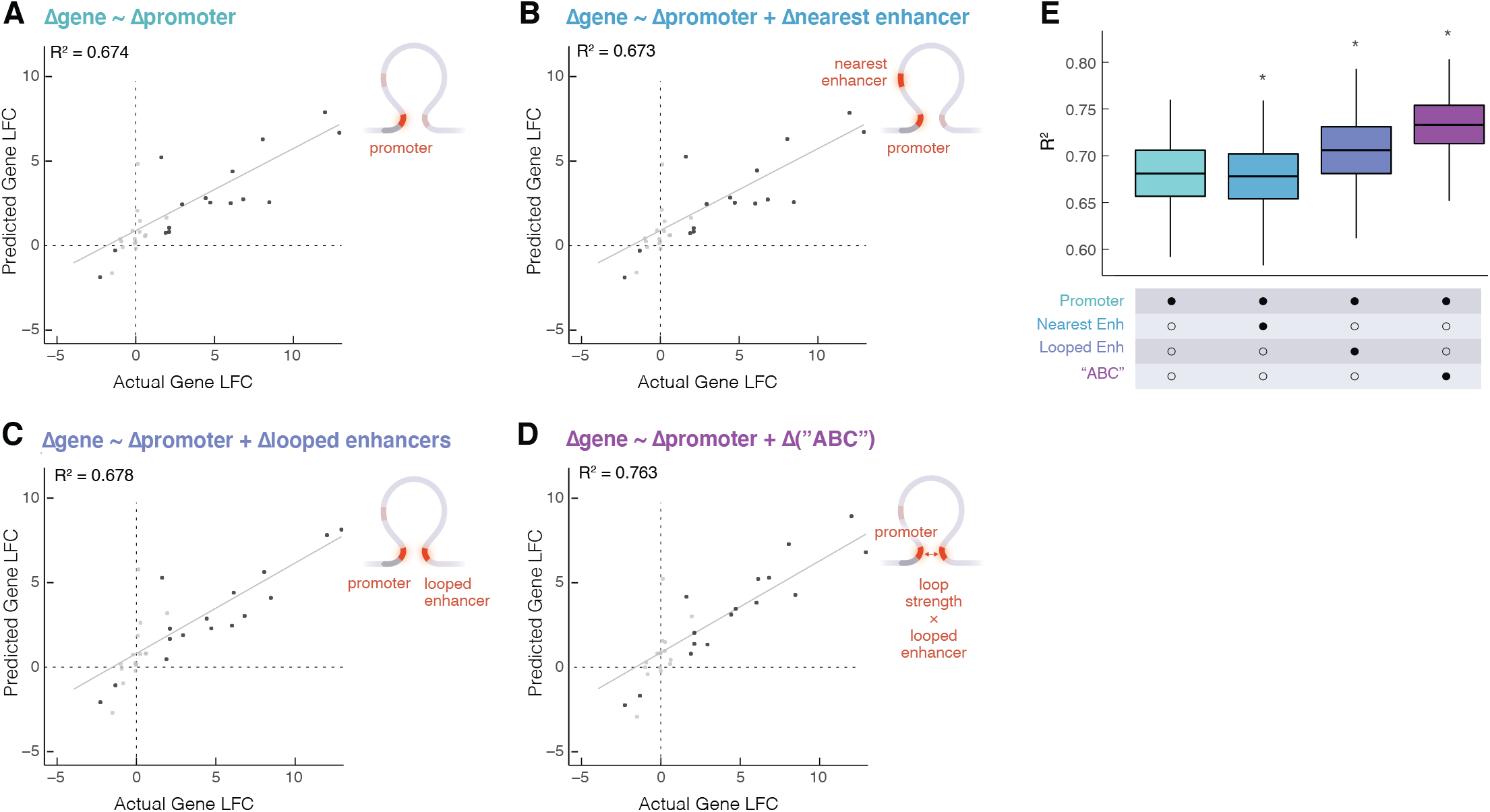
Changes in gene expression at differential loops are explained by combined proximal and distal enhancer activity and loop strength. Scatter plots showing predicted gene-fold change vs actual gene fold-change for genes that are at the anchors of differential loops based on **(A)** promoter H3K27 ac LFC alone, **(B)** promoter H3K27ac LFC and the nearest enhancer to the promoter’s FC, **(C)** promoter H3K27ac FC and distal looped H3K27ac, and **(D)** promoter H3K27ac LFC and the LFC of the product of distal looped H3K27ac and loop strength (red = differential gene, gray = static gene). **(E)** R2 for each model calculated based on 1000 permutations of splitting data into training and testing sets. Wilcoxon rank sum test was performed to compare each group to the promoter only model, *asterisk represents p < 0.05.

